# Komagataeibacter tool kit (KTK): a modular cloning system for multigene constructs and programmed protein secretion from cellulose producing bacteria

**DOI:** 10.1101/2021.06.09.447691

**Authors:** Vivianne J Goosens, Kenneth T Walker, Silvia M Aragon, Amritpal Singh, Vivek R Senthivel, Linda Dekker, Joaquin Caro-Astorga, Marianne L A Buat, Wenzhe Song, Koon-Yang Lee, Tom Ellis

## Abstract

Bacteria proficient at producing cellulose are an attractive synthetic biology host for the emerging field of Engineered Living Materials (ELMs). Species from the Komagataeibacter genus produce high yields of pure cellulose materials in a short time with minimal resources, and pioneering work has shown that genetic engineering in these strains is possible and can be used to modify the material and its production. To accelerate synthetic biology progress in these bacteria, we introduce here the Komagataeibacter tool kit (KTK), a standardised modular cloning system based on Golden Gate DNA assembly that allows DNA parts to be combined to build complex multigene constructs expressed in bacteria from plasmids. Working in *Komagataeibacter rhaeticus*, we describe basic parts for this system, including promoters, fusion tags and reporter proteins, before showcasing how the assembly system enables more complex designs. Specifically, we use KTK cloning to reformat the *Escherichia coli* curli amyloid fibre system for functional expression in *K. rhaeticus*, and go on to modify it as a system for programming protein secretion from the cellulose producing bacteria. With this toolkit, we aim to accelerate modular synthetic biology in these bacteria, and enable more rapid progress in the emerging ELMs community.

## Introduction

The emerging field of Engineered Living Materials (ELMs) uses synthetic biology to grow and engineer materials with characteristics that are prominent in nature, such as colour, self-repair, growth, conductivity and inherent sensing (Gilbert and Ellis, 2019; Tang et al., 2020). Being biological, these materials also remain biodegradable and can be grown from sustainable nutrient sources and so have great potential in a circular economy. A significant focus of early ELMs work has been on bacterial biofilms, and in particular, on manipulating proteinaceous amyloid fibres and cellulosic polymers as these lends themselves to a number of ELM-based applications (Gilbert and Ellis, 2019; Nguyen et al., 2018).

Bacterial cellulose (BC) is a a major component of biofilms and is unique in its purity, synthesis and network architecture, offering a biocompatible material with superior crystallinity and tensile strength compared to plant cellulose (Pallach et al., 2018; Yadav et al., 2010). These physiochemical characteristics of BC has made it valued in the food, beverage, cosmetic, surgical and biomedical industries and has enormous potential as an ELM (Gilbert and Ellis, 2019; Jang et al., 2017). Gram-negative acetic acid bacteria (AAB), in particular Gluconacetobacter and Komagataeibacter species, are prolific producers of BC.

In recent years, many advances have been made in molecular tools for the BC field (Florea et al., 2016; Gwon et al., 2019; Singh et al., 2020; Teh et al., 2019; Walker et al., 2018), however, both the general cellulose and the more specific cellulose-based ELM fields could greatly benefit from a Golden Gate (GG) cloning toolkit. Modular GG-based toolkits have revolutionised molecular and synthetic biology in other organisms including in yeasts, mammalian cells, plants, and gram-positive and gram–negative bacteria (Andreou and Nakayama, 2018; Engler et al., 2014; Hernanz-Koers et al., 2018; Iverson et al., 2016; Lee et al., 2015; Martella et al., 2017; Moore et al., 2016; Weber et al., 2011; Wicke et al., 2017). Toolkits with both broad or specific host-ranges, have been developed and are transforming their relevant fields (Chiasson et al., 2019; Geddes et al., 2019; Valenzuela-Ortega and French, 2019; Vasudevan et al., 2019; Wu et al., 2018). These cloning systems allow for rapid assembly of DNA constructs from modular genetic parts to build multigene systems. This is achieved using specially-designed complementary overhangs that are created by Type IIS restriction enzymes and joined by ligases.

Recognising a lack of such a system for BC-producing bacteria, we developed and herein describe the Komagataeibacter Tool Kit (KTK): a modular cloning system that caters to Komagataeibacter species, the BC biosynthesis field and those developing cellulose-based ELMs. We define our KTK system and the plasmids and part standards associated with it, and then demonstrate its use in *Komagataeibacter rhaeticus*, a transformable native BC producer, in cases using basic parts for gene expression, and then for cases where multigene expression is necessary to achieve new functionality. As a case-study, we used KTK cloning to express *E. coli*’s Curli system, which produces the best-studied bacterial amyloid; a polymeric protein structurally rich in beta-sheets that forms a fibre with exceptional strength, stability and resistance (Evans and Chapman, 2014). Curli has been a focus for many ELM-associated studies (Birnbaum et al., 2020; Chen et al., 2014; Nguyen et al., 2014; Seker et al., 2017), and offers promise for making a novel composite with BC in cells that can co-produce both materials. The Curli production system is also representative of a Type VIII secretion system (T8SS) and so its expression offers a tractable solution for protein secretion from BC-producing cells. Expressing this system in *K. rhaeticus* allows us to secrete Curli proteins and enzymes from BC-producing bacteria and demonstates the power of the KTK system for assembling complex multigene modular assemblies.

## Results

### The Komagataeibacter Toolkit (KTK)

The KTK system is an iterative, hierarchical Golden Gate (GG) cloning method best described in levels (**Fig 1a**). GG cloning takes advantage of Type IIS restriction enzymes (REs). These enzymes cut a short distance away from the recognition site to generate specific overhangs used for orientated ligations. Once these overhangs are joined, the DNA fragment no longer includes the initial RE site, thereby cementing the ligation and prevent re-cutting (for detailed guide and illustrations see **Supplementary Materials**). This GG system starts with *Entry-level Parts*, the basic DNA-encoded components required for gene expression in a single Transcriptional Unit (TU). Entry level parts include promoters (E1.1), a ribosome-binding site (RBS) sequences (E1.2), coding sequence (CDS) regions (E1.3) and terminators (E1.4) (**Fig 1b**). The Entry-level Parts are designed to include complementary GG overhangs for oriented ligations, so that when combined in the first level of cloning with a *Destination* vector backbone they ligate to form a plasmid with a single full Transcription Unit (**Fig 1b**).

**Figure 1.**
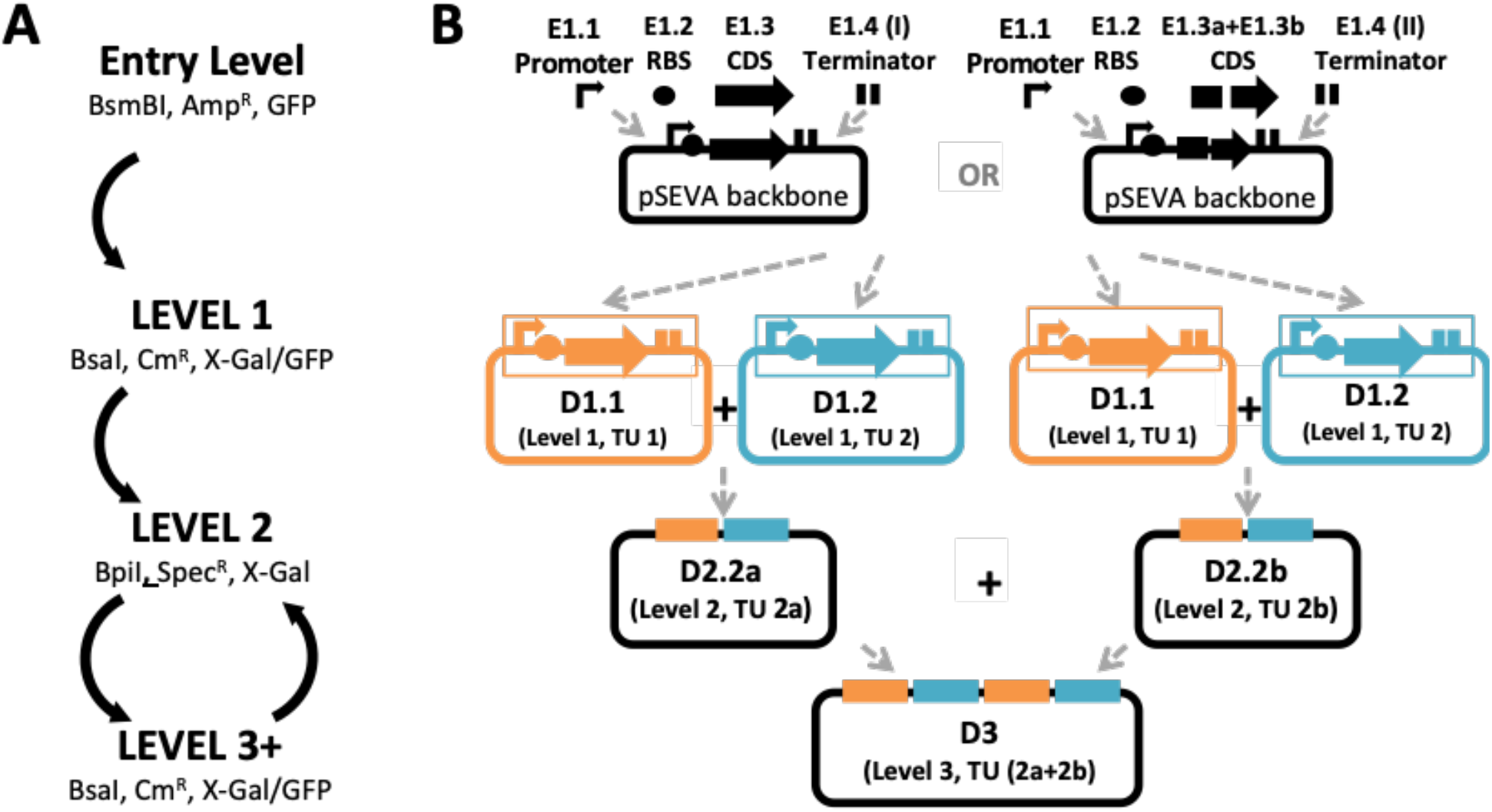
The KTK cloning system. **(a)** The levels of KTK cloning. Entry-level plasmids are ampicillin resistant (Amp^R^), are selected by the absence of GFP, and BsmBI is used for cloning. In Level 1 (and Level 3) cloning plasmids generated using BsaI, resulting in chloramphenicol resistant (Cm^R^) plasmids that have lost the X-Gal or GFP marker cassette. BpiI is used in Level 2 cloning and resultant plasmids are spectinomycin resistant and selected for by loss of X-Gal cassette. **(b)** Schematic of the Entry-level parts combining with a Level 1 backbone vector to form either a basic TU (made up of promoter (E1.1), RBS (E1.2) CDS (E1.3) and terminator (E1.4I) parts or a fusion protein TU (made up of promoter (E1.1), RBS (E1.2), fused CDS (E1.3a +E1.3b) and terminator (E1.4II) parts. There are two variations of Level 1 destination vectors and two variation of Level 2 destination vectors. Combined this allows the combination of 4 TUs at the 3^rd^ cloning level. In both Level 1 and Level 2 cloning there are two possible destination vector backbones (D1.1 & D1.2 and D2.2a & D2.2b, respectively). The 3^rd^ cloning level recycles the same destination vectors as in Level 1, facilitating iterative cycles of cloning between the two vector backbones.

KTK Destination Vectors are based on Standard European Vector Architecture (SEVA) plasmids, and benefit from the advantages of this powerful modular set (Durante-Rodríguez et al., 2014). The main destination vectors are adapted from the pSEVA331 and pSEVA431 plasmids, which have a pBBR1 replication origin and encode Chloramphenicol and Spectinomycin resistance markers, respectively. In the KTK system, the multiple cloning site (MCS) regions of these plasmids has been modified to encode Golden Gate cloning sequences. A further possibility when constructing a TU with KTK is to create a fusion protein in the CDS position (E1.3). The system allows this by ligating C- and N-terminal domain CDS parts (E1.3a and E1.3b) (**Fig 1b**). The basic KTK system includes a library of useful Entry-level Parts (promoters, RBS, terminators, and useful C- and N-terminal CDS domains e.g. GFP, His-tag and signal peptides) as well as sequences that can be used as ‘spacers’ that are useful at subsequent cloning levels. All plasmid construction is designed to be done using *E. coli* cloning strains, before completed plasmids are then transformed into competent Komagataeibacter for testing.

For simple constitutive expression of a protein, constructing a TU via a single round of cloning (level one) is sufficient. However, for multifaceted genetic constructs, such as inducible expression cassettes or multi-part TUs, the KTK adds further cloning levels that enable versatile construction. The backbone vectors for each cloning level have an alternating arrangement of flanking Type IIS RE sites (for BsaI and BpiI). These in turn generate specific overhangs that are used for the next level of cloning (illustrated and detailed in **Supplementary Materials**). The nature of Type IIS RE cloning means that one of set of RE sites are removed during the ligation reaction and the second set then becomes available for the next level of assembly. Furthermore, the KTK system provides two Backbone vectors for each cloning level, each with slight variation in overhangs to allow for the parallel insertion of different ligated DNA parts and TU assemblies. These can then be joined in the next level of ligation, thereby facilitating multiple cycles with multipart constructs (**Fig 1b**).

### Validating KTK Golden Gate cloning

To validate Level 1 Golden Gate assembly with KTK, plasmids were constructed to express fluorescent proteins from encoded TUs. Entry-level Parts were first prepared, including the well-characterised constitutive promoter J23104 (E1.1) (Florea et al., 2016; Walker et al., 2018), a standard RBS (E1.2), a terminator (E1.4) and CDS parts (E1.3) encoding two fluorescent proteins, superfolder Green Fluorescent Protein (sfGFP) and mScarlet Red Fluorescent Protein (RFP). These two CDS parts were also cloned as N-terminal (E1.3a) and C-terminal (E1.3b) CDS parts to also allow assembly of a TU expressing sfGFP-mScarlet fusion protein (**Fig 2a**). Assembly of basic and fusion TU constructs was done using the KTK system with cloning steps in *E. coli*. The three constructs were then transformed into *K. rhaeticus* competent cells, and selected transformant colonies were then cultured in liquid growth media in the presence of purified cellulase (to prevent material formation) and measured for green and red fluorescence by flow cytometry (**Fig 2b**). The fluorescence intensity per cell compared to untransformed cells showed strong expression of both reporter proteins in basic TU form, and expression with reduced strength when constructed as a fusion protein.

**Figure 2.**
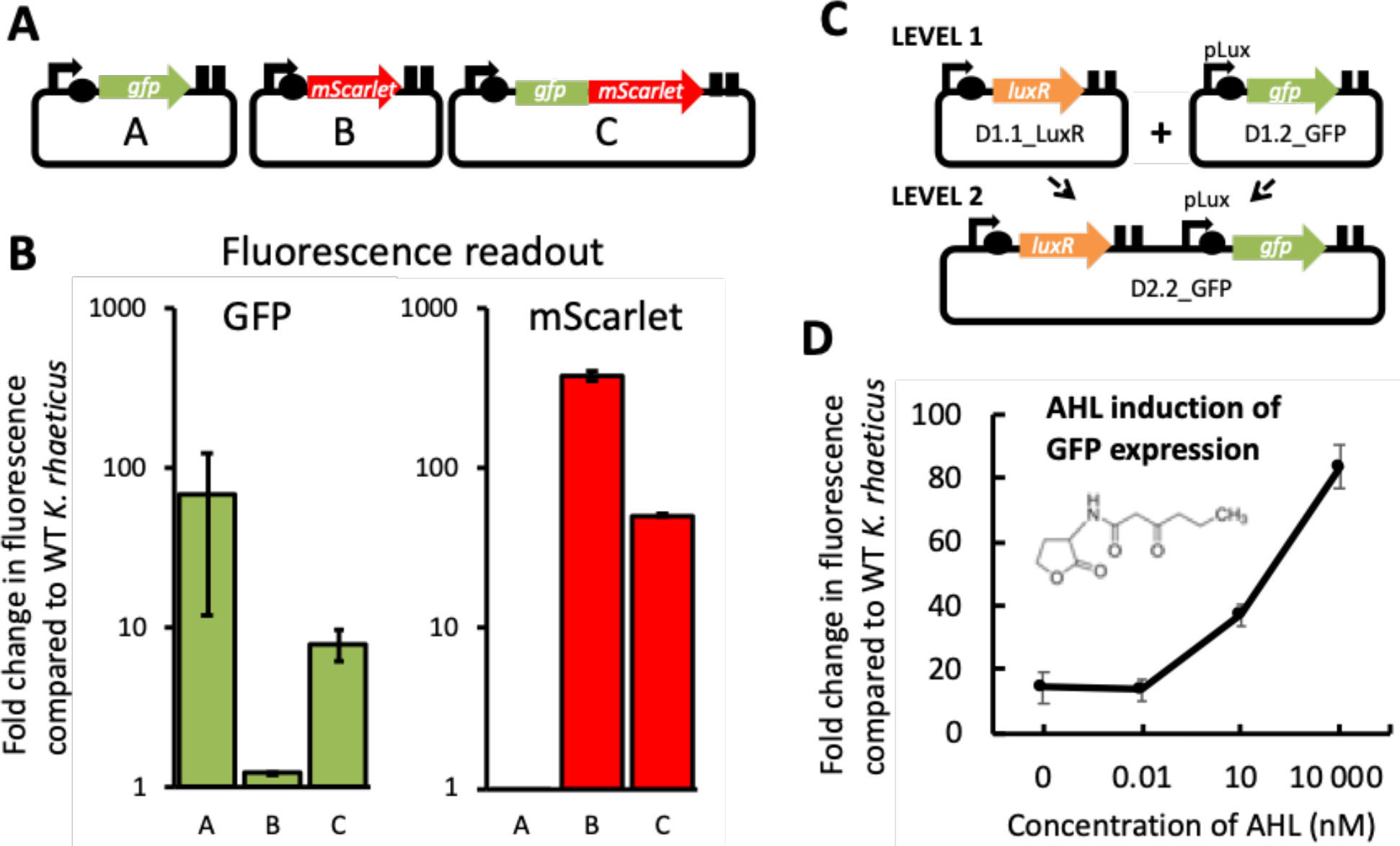
Validation of KTK cloning for basic TU and fused TU constructs (Level 1) and a multigene cassette (Level 2). **(a)** Design of the assembled fluorescent protein expressing constructs used to validate KTK level 1 cloning. All promoters used are J23104. **(b)** Fluorescence per cell measured by flow cytometry of *K. rhaeticus* strains expressing GFP, mScarlet or a fused GFP-mScarlet protein. Values normalised to untransformed *K. rhaeticus* with GFP fluorescence shown in green and mScarlet in red. Error bars represent SD of three replicates. **(c)** Cloning schematic showing the use of different Destination vectors to enable Level 1 to Level 2 cloning. Level 1 assembly results in a LuxR-expressing TU (with J23104 promoter), and a TU with pLux promoter and GFP. Level 2 cloning brings these TUs togethers to give an AHL-inducible GFP expressing plasmid. **(d)** *K. rhaeticus* strains with the plasmid constructed in (c) were exposed to a range of exogenous AHL concentrations in liquid growth phase and GFP fluorescence per cell was measured by flow cytometry. Error bars represent SD of three replicates.

Next, a multigene Level 2 construct was designed to yield a plasmid encoding externally-inducible GFP expression. Previous *K. rhaeticus* studies have taken advantage of the Vibrio Lux quorum sensing system for external gene expression induction (Florea et al., 2016; Teh et al., 2019; Walker et al., 2018). In the Lux system a regulator protein (LuxR) binds and induces expression of the gene coupled the pLux promoter when LuxR is bound to an acyl homoserine lactone (AHL). In the design chosen here, LuxR-AHL triggers expression of GFP. This two-gene construct was assembled by KTK cloning from Entry-level parts. Unlike in cloning of a single TU, choice of Level 1 Destination vectors is important when going to Level 2. A TU constitutively expressing LuxR was assembled into a D1.1 vector, while the TU expressing sfGFP from the pLux promoter was assembled into a D1.2 vector (**Fig 2c**). The multigene Level 2 construct was then assembled from Level 1 constructs and transformed into *K. rhaeticus*. This construct worked as expected showing induction of GFP expression in the presence of increasing AHL concentrations (**Fig 2d**).

### Expanding and characterising the KTK parts library

The utility of GG-based cloning toolkits are dependent on the size, characterization and availability its Parts. To populate the KTK system we prepared a library of shareable Entry-Level Parts as well as useful Destination-Level plasmids (**Table S1**). This library includes a panel of promoters, RBS sequences, terminators, and basic CDS parts (selection cassettes, fluorescent proteins) as well as spacer sequences that enable more complex assembly steps. Our basic library will be made available to researchers and is intended to become a resource for the wider community.

As tuning gene expression is a key goal in engineering cells, we characterised a panel of promoters including heterologous promoters and promoters native to *K. rhaeticus,* with the latter identified from genes and operons known to express from the strain genome. KTK enabled straightforward and quick assembly of modular constructs expressing sfGFP in basic TUs, each with a different promoter. These promoters were then characterised for expression strength in liquid growth phase using flow cytometry quantification of green fluorescence. J23104 and pTac promoters were the strongest heterologous promoters (**Fig 3a**), while the native promoters pTtcA and pHxu were the strongest of the ones taken from the genome sequence (**Fig 3b**). The individual strengths of the heterologous promoters in KTK format correlate well to their strengths measured in past work in this strain in different DNA formats (Florea et al., 2016; Teh et al., 2019).

**Figure 3.**
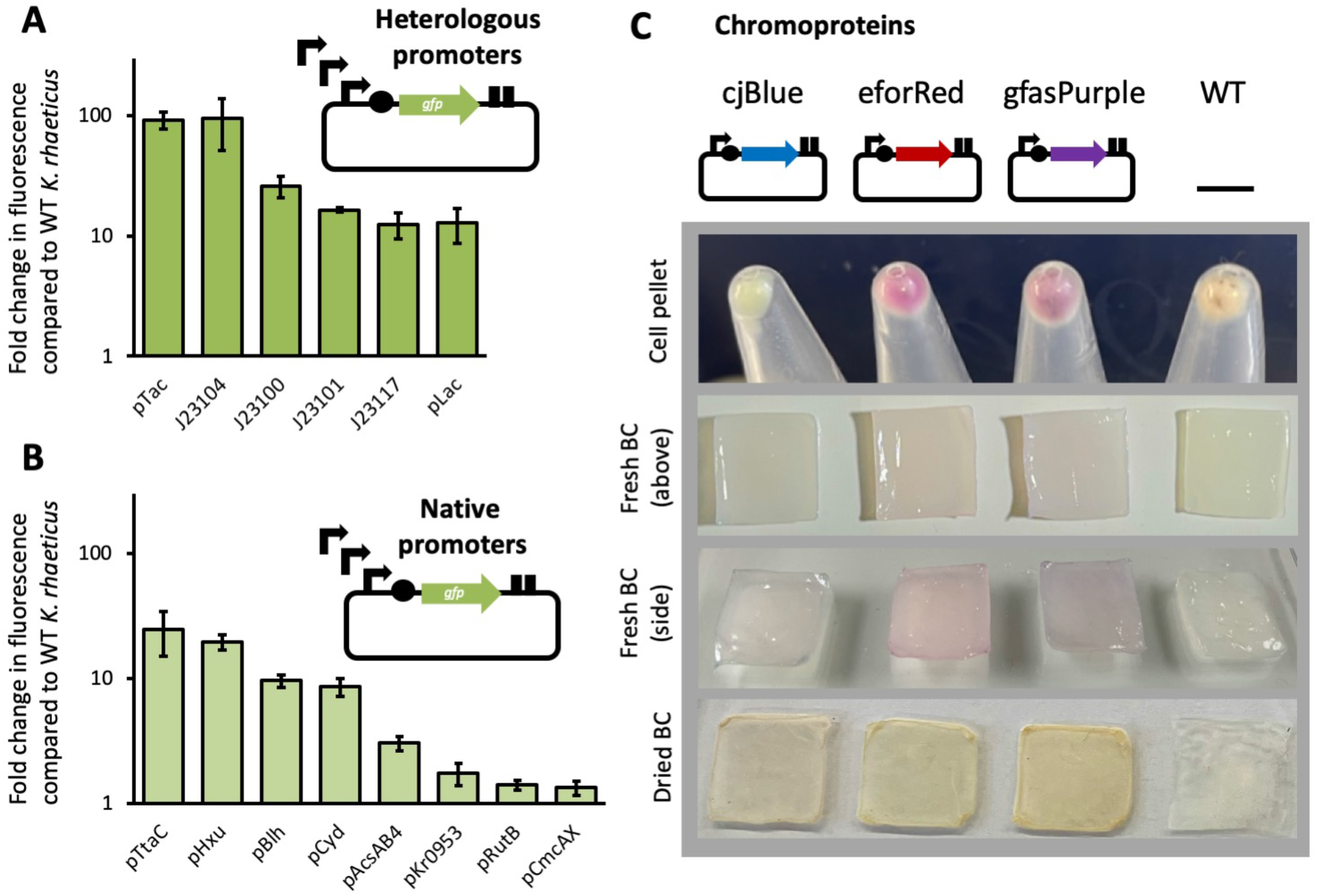
Characterization of promoters for the KTK parts library. **(a)** Fluorescence per cell measured by flow cytometry of *K. rhaeticus* strains expressing GFP in a standard single TU from 6 different heterologous promoters. **(b)** Fluorescence per cell measured by flow cytometry of *K. rhaeticus* strains expressing GFP in a standard single TU from 8 native promoters. Values in the plots are normalised to untransformed *K. rhaeticus*. Error bars represent SD of three replicates. **(c)** Cell pellets (top row), fresh BC pellicles (middle two rows) and air-dried BC pellicles (bottom) from cells transformed with plasmids expressing 3 representative chromoprotein reporters.

We further sought to add reporter proteins visible to the naked eye that can be expressed within cellulose materials. To this end, we cloned 3 chromoproteins previously characterised in *E. coli* (Liljeruhm et al., 2018) that are visibly blue (cjBlue), red (eforRed) and purple (gfasPurple). When expressed in *K. rhaeticus* as a level 1 construct with J23104 promoter, each chromoprotein was able to change the colour of the bacterial cell pellet (**Fig 3b**), although cjBlue gave a more greenish hue than expected. The resulting bacterial cellulose pellicles grown from these strains showed visible colouration of the material, with the red and purple colours being particularly striking. However, on dehydration of the cellulose, pigmentation was lost, although the final material was still notably a different shade to cellulose made from unmodified cells (**Fig 3b, Fig S3**).

The basic KTK parts library contains several reporter proteins, terminators, RBS sequences and constitutive promoters, as well as the AHL-inducible promoter (**Fig 2**). We aim to expand this soon to include more inducible expression systems and constructs that allow for CRISPR-based gene regulation, and share the most relevant plasmids as a distributable collection.

### KTK construction of an inducible curli system

For a case-study application of the KTK system, we next designed and constructed a multigene assembly to express and extrude curli amyloid fibres from *K. rhaeticus*. In *E. coli*, curli fibres are polymerised on the outer surface of the cell from monomers of the CsgA protein, with this facilitated by five other curli proteins that act to chaperone (CsgC) and nucleate (CsgB) fibre formation, and form a curli-specific type VIII secretion system (T8SS) (CsgE, CsgF, CsgG) through the outer membrane (Evans and Chapman, 2014; Van Gerven et al., 2015) (**Fig 4A**). In *E. coli* these genes are encoded on two divergent operons, however a number of landmark ELM studies have elegantly functionalised curli by rearranging these genes into an easier-to-engineer synthetic linear operon (Kan et al., 2019). The linear operon format enables researchers to more readily modify curli through C-terminal CsgA fusions (small peptides or whole proteins) or by exploiting the signal peptide (the first 22 amino acids of CsgA) to secrete heterologous proteins through the Curli T8SS (Assalkhou et al., 2007; Birnbaum et al., 2020; Botyanszki et al., 2015; Nguyen et al., 2014; Nussbaumer et al., 2017; Seker et al., 2017; Sivanathan and Hochschild, 2012; Tay et al., 2017; Van Gerven et al., 2014; Wang et al., 2017; Zhong et al., 2014).

**Figure 4.**
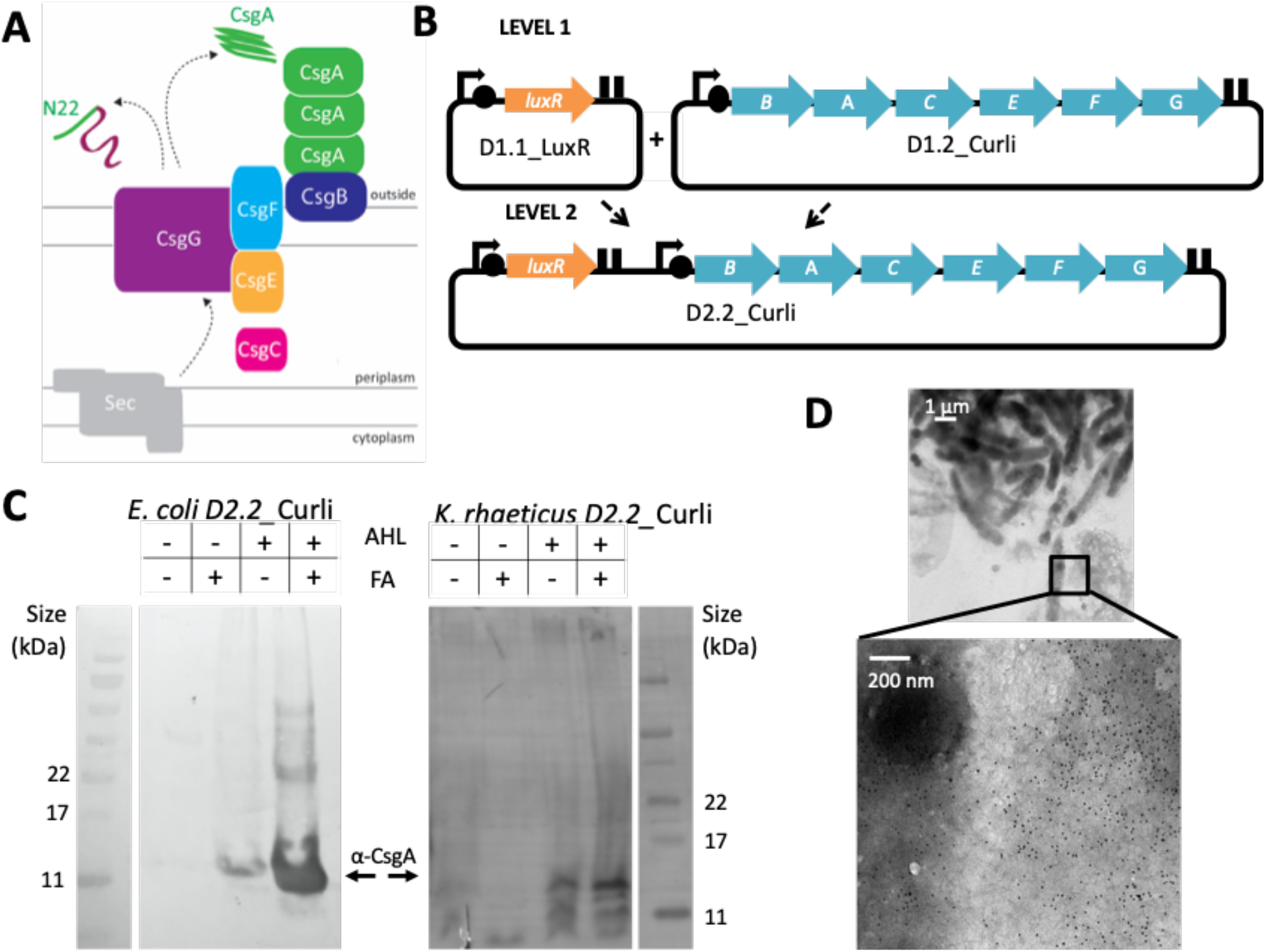
Expressing an inducible Curli system in *K. rhaeticus*. **(a)** Schematic of the *E. coli* curli system where amyloid fibres are assembled composed of polymers of CsgA, nucleated to the cell surface via CsgB. These proteins are chaperoned (CsgC) and transferred over the outer membrane via a specialised Type VIII secretion (T8S) system (CsgE, CsgF, CsgG). All proteins are secreted into the periplasm via the Sec system, however, the N22 region of CsgA (shown in green) is essential for directing the protein through the CsgG pore in the outer membrane. **(b)** Cloning schematic illustrating the KTK assembly of the AHL-inducible, Curli-expressing D2.2_Curli plasmid from two level 1 plasmids. **(c)** Western blot of protein extractions from *E. coli* and *K. rhaeticus* cultures transformed with the D2.2_Curli plasmids. Cultures were grown with or without AHL inducer and obtained extracts were treated with or without Formic Acid (FA) to denature Curli proteins. Polyclonal anti-CsgA antibody was used for detection and the CsgA band is indicated with an arrow. **(d)** Transmission electron microscopy visualisation of gold particle-stained curli proteins outside of a D2.2_Curli expressing *K. rhaeticus* (inset). Black dots in the high magnification image are gold nanoparticles bound to the curli protein via α-CsgA rabbit antibody and 10 nm anti-rabbit IgG-Gold coupled antibody.

To port the *E. coli* curli system into BC-producing bacteria, we chose to maintain the linear operon format successfully employed in past ELMs work. The region encoding the 6 genes from the synthetic *E. coli* operon was therefore cloned into a KTK Entry-level vector as if it was a single CDS (E1.3) part (**Fig 4B**). This was then assembled into a Level 1 curli-containing plasmid (D1.2_Curli), with the AHL-inducible pLux promoter (E1.1), a well-characterised terminator (E1.4I) and an RBS (E1.2) part, selected by the RBS Calculator (Salis et al., 2009) to give a similar predicted strength for CsgB translation as that seen in the native *E. coli* system. D1.2_Curli was then assembled with a Level 1 LuxR-expressing plasmid (D1.1_LuxR) to create the multigene Level 2 plasmid construct giving AHL-inducible Curli expression (D2.2_Curli). In this design LuxR is expressed by a medium strength promoter (pLac) to reduce any potential burden.

As KTK constructs are compatible in *E. coli,* assays for plasmid function can be performed in *K. rhaeticus* and *E. coli* in parallel. To test for expression and production of CsgA from the D2.2_Curli construct, we first used Western blot analysis. As amyloid fibres are resistant to heat and detergents, formic acid (FA) treatment is required for depolymerisation SDS-PAGE separation of CsgA and so can be used to identify if the CsgA is polymerised or just in monomeric form. Protein extracts taken from uninduced or AHL-induced *K. rhaeticus* and *E. coli* were separated and stained (**Fig 4c**) for monomeric CsgA (visible band with no FA treatment) and polymeric CsgA (visible band only seen with FA treatment), confirming that the D2.2_Curli construct produced CsgA and polymerised it into curli fibres in both *K. rhaeticus* and *E. coli*.

Transmission electron microscopy (TEM) was next performed to visualise curli fibre production from transformed *K. rhaeticus.* As control, TEM analysis of AHL-induced *E. coli* showed clear filamentous Curli structures (**Fig S1a**) confirming the plasmid function in this bacterium. Direct visualisation was more challenging with *K. rhaeticus* due to cellulose fibres being present in large amounts, even after 20% (w/v) cellulase treatment (**Fig S1b**). Therefore, to distinguished curli from cellulose, TEM samples were treated with CsgA-polyclonal antibody and immunogold-labelled. Gold nanoparticle staining was evident in a tangle of fibres outside the *K. rhaeticus* cell, demonstrating that the plasmid is functional in the cellulose-producing bacterium (**Fig 4D and Fig S1b**). The effect of induced co-production of curli on the material properties of grown bacterial cellulose was then investigated to see if this cellulose-amyloid composite had greater strength than cellulose alone. Strength tests were performed on pellicles grown from AHL-induced *K. rhaeticus* with the D2.2_Curli construct (**Fig S1c**). Pellicles grown from cells containing the D2.2_GFP construct were used as a comparable control. No significant difference in strength was observed between the control and curli-positive pellicles, likely due to a very low relative abundance of curli compared to cellulose in the final material.

### Programming protein secretion via the Type VIII secretion system

For an example of a Level 3 KTK multigene assembly, we next constructed a plasmid that exploits the curli-specific T8SS system for heterologous protein secretion (**Fig 5a**). Previous studies have shown that the first 22 amino acids (N22) of CsgA are enough to target small heterologous proteins for secretion via T8SS (Sivanathan and Hochschild, 2012; Wang et al., 2017). To make use of this we split the curli operon into modules, with the 3 genes encoding the T8SS (CsgE, CsgF, CsgG) as a key part and the N22 signal used as a tag to be fused to proteins for secretion (**Fig 5a**).

**Figure 5.**
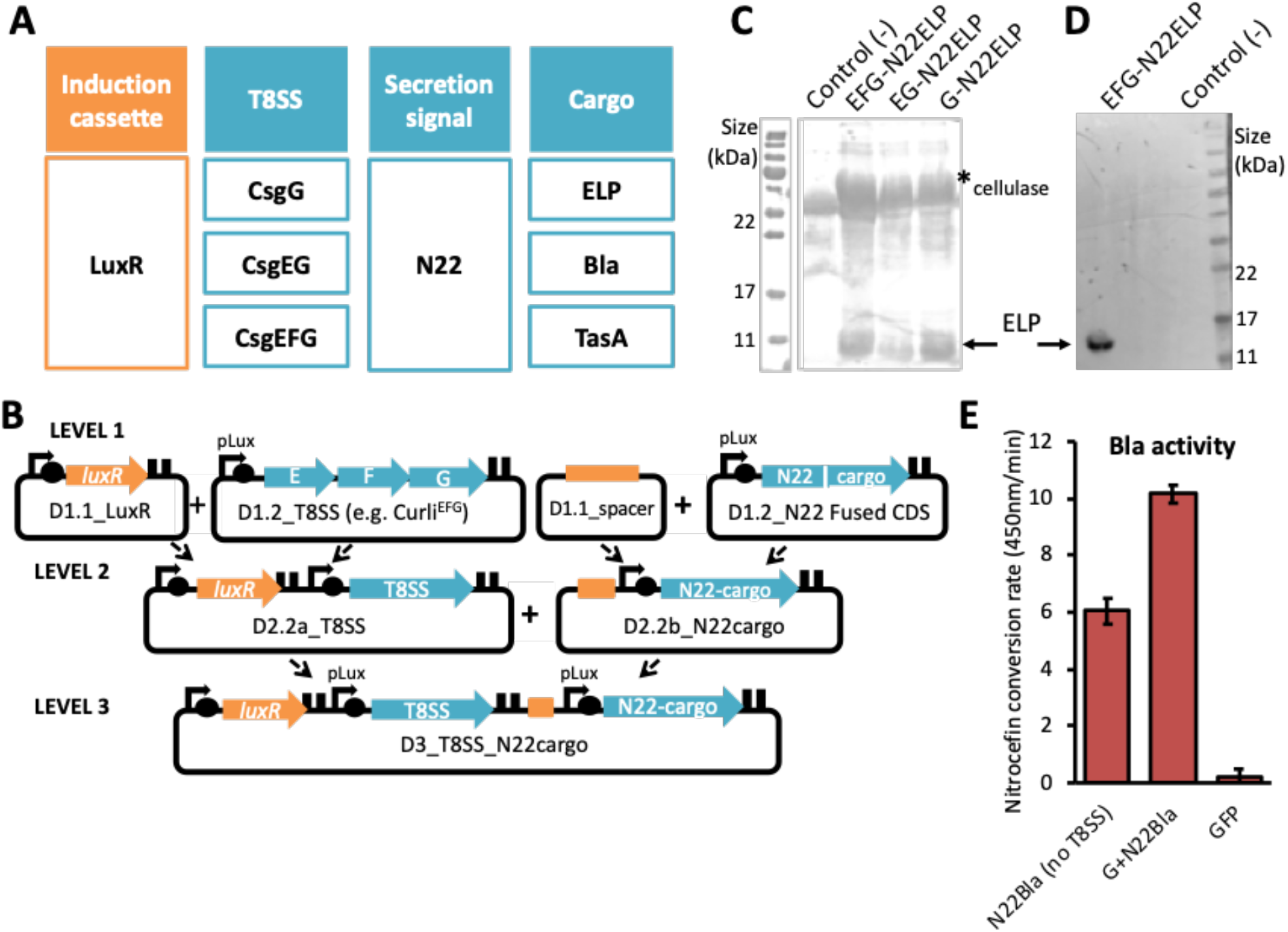
Using a T8SS module to secrete heterologous cargo proteins. **(a)** Modular components of the KTK-based T8SS constructs. The first module (orange) represents the induction cassette (LuxR) for controlling gene expression, further modules (teal) are 3 versions of the T8SS (CsgG, CsgEG and CsgEFG), the secretion signal (N22) and the heterologous cargo proteins (ELP, Bla and TasA). **(b)** Cloning schematic to generate an AHL-inducible modularised T8SS for heterologous protein secretion using the KTK system. Four level 1 constructs are made: an induction cassette construct (D1.1), the secretion system construct (D1.2), a spacer construct (D1.1) and constructs that fuse the secretion signal (N22 sequence) to the cargo protein (D1.2). These are combined in pairs to level 2 into the D2.2a and D2.2b vectors. A final assembly creates the Lebel 3 construct, D3_T8SS_N22cargo, which contains both an inducible secretion target and secretion system. **(c)** SDS-PAGE (left) and Western Blot (right) of samples extracted from culture supernatants from AHL-induced *K. rhaeticus* strains expressing a T8SS module (CsgEFG, CsgEG and CsgG) and N22-ELP-his. To aid liquid phase growth and remove cellulose material, exogenous purified cellulase was added to the cultures and is clearly visible in the SDS-PAGE protein gel. **(d)** Nitrocefin assay measuring the Beta-lactamase (Bla) activity of supernatant harvested from an induced *K. rhaeticus* strain designed to express CsgG and N22-Bla. An inducible strain expressing GFP (D2.2_GFP) was used as a negative control in this experiment. Error bars represent SD of three replicates.

A panel of three T8SS modules were cloned into KTK as CDS parts; the minimal T8SS (just the secretion pore CsgG), CsgG plus the CsgE ‘adapter protein’ (CsgGE), and both of these proteins plus secretion chaperon CsgF (CsgEFG) (Costa et al., 2015; Hospenthal et al., 2017). These were assembled to Level 2 with a LuxR TU, so that expression of these T8SS modules was AHL-dependent (**Fig 5b**). In parallel, a small panel of heterologous cargo targets were prepared using the 2-part CDS fusion approach of KTK to bring together the N-terminal N22 region of CsgA (E1.3a part) with C-terminal CDS parts (E1.3b) encoding three targets: a 98 amino acid elastin-like protein (ELP), the enzyme Beta-lactamase (Bla) and the Gram-positive amyloid protein TasA (**Fig 5a-b**). ELPs are small unstructured versatile proteins that are highly hydrophobic and an attractive ELM component (Roberts et al., 2015), Beta-lactamase is a classic enzyme easily detected by the colorimetic nitrocefin assay (Gilbert et al., 2021), and TasA is an amyloid protein from *B. subtilis* that could be an alternative to curli (Erskine et al., 2018).

The 3 heterologous cargo targets were assembled into a Level 1 TU construct with the pLux promoter, before then being each combined into a Level 2 construct with a short spacer part (D1.2_Spacer, 23 bp) designed to assist cloning of an odd number of TUs in a multigene assembly (**Fig 5b**). With all modules now in place in Level 2 plasmids, a combinatorial set of nine possible Level 3 constructs could be assembled with the 3 different T8SS versions and 3 cargo proteins (**Fig 5a**).

To test the function of these constructs, we again performed experiments in parallel in *E. coli* and *K. rhaeticus* transformed with constructs, culturing with AHL induction and then harvesting in mid-log growth phase and separating supernatant and cell fractions. We first assessed ELP expression and secretion, using SDS-PAGE of both *K. rhaeticus* (**Fig 5c**) and *E. coli* (**Fig S2a**) samples to confirm the presence of processed and secreted ELP when any one of three T8SS versions were present. As ELP proteins are difficult to transfer by Western Blot due to their hydrophobic nature, the constructs were designed to include a C-terminal his-tag. This enabled us to identify that the 10 kDa protein observed in the secreted fraction was indeed ELP-his when either *K. rhaeticus* (**Fig 5d**) or *E. coli* (**Fig S2b**) were expressing the CsgEFG T8SS constructs. Similar experiments with the TasA cargo were not as successful, only showing a small amount of TasA in cell fractions of *E. coli* (**Fig S2c**), suggesting that TasA cannot use the curli T8SS for export.

Finally, we assessed enzyme secretion from the beta-lactamase (Bla) encoding Level 3 constructs, as before testing in both *E. coli* and *K. rhaeticus*. Beta-lactamase converts nitrocefin from a yellow to red colour, which is quantifiable by spectroscopy at 450nm and initial *E. coli* data showed that in the presence of either T8SS version, Bla enzymatic activity in the supernatant was higher than when no T8SS was present (**Fig. S2d**). In this case the minimal T8SS (CsgG) gave the highest values, so the Level 3 construct expressing this with N22-Blac was then assessed in *K. rhaeticus*. Supernatant from an AHL-induced culture of these bacteria showed beta-lactamase activity nearly 2-fold higher than when no T8SS is expressed in the cells (**Fig 5d**). Altogether the results with ELP and Bla cargos show that the modular T8SS system generated here for the KTK system can be used to program protein secretion from K. rhaeticus. Further work is needed to understand the optimal T8SS version, its ideal expression level and what design considerations are needed for protein cargos and their secretion.

## Conclusion

Through a standardised hierarchical Golden Gate DNA assembly approach, the KTK system allows modular construction of single genes and multigene systems for cellulose-producing Komagataeibacter. We here validated this toolkit using fluorescent protein and chromoprotein constructs, before applying it to the modular construction of plasmids that enable inducible production of curli fires by *K. rhaeticus* and heterologous protein secretion by this strain. These examples demonstrate the power of the combinatorial and modular toolkit system.

We provide a selection of basic parts along with the KTK system for tuning, controlling and measuring gene expression. Using promoters and RBS parts of different strength is a powerful way to optimise gene function in bacteria. Most of the experimental results shown here were done without any optimisation of expression of assembled genes. Design-led or combinatorial approaches choosing from promoter and RBS libraries are likely to be able to improve on the results seen here for the various demonstrations, for example in helping to make an updated GFP-RFP fusion construct (**Fig 2c**) that maintains equivalent fluorescence to constructs expressing only one of the two proteins.

To demonstrate how multigene assembly with the KTK system can advance engineered living materials, we imported the well-studied *E. coli* curli fibre production system into KTK and used it to engineer a construct that instructs *K. rhaeticus to* co-secrete curli fibres alongside cellulose as material is produced. Although KTK-based cloning was successful in enabling the production of amyloid curli outside the cell, this did not result in any marked change in material properties for the cellulose-curli composite pellicle. This is likely due to relatively very low production of protein compared to cellulose from these bacetria, or could be related to curli fibre formation being impaired chemically by the low pH of *K. rhaeticus* cultures, or physically by the large amounts of cellulose being extruded from the cell surface. In future work, curli production could be improved by increasing gene expression with different promoter/RBS parts and by using alternative gene arrangements, *e.g.* by using strong promoters and a two-module design (Bongers et al., 2005; Mierau et al., 2005; Tabor, 1990).

The potential of the curli system was further exploited here by taking advantage of its Type VII secretion system (T8SS) to export heterologous cargo proteins. An N-terminal fused N22 region from CsgA enabled secretion of a small unstructured ELP proteins when expressed in cells co-expressing the CsgEFG T8SS. The tag also gave promising results with Bla for the secretion of active globular enzymes from *K. rhaeticus* strains expressing CsgG. This offers the first described route to getting enzymes expressed in BC-producing bacteria to be secreted extracellularly, opening up the possibility of having cellulose-binding or cellulose-modifying enzymes co-produced as the cellulose material is grown. However, the relatively low levels of protein and enzyme activity observed, and the failure to secrete TasA here suggests that achieving high-level programmed secretion of a desired target protein will more often than not be a challenge and will likely to require significant optimisation. We hope that many people will use and contribute to the KTK system in the future, to add more genetic parts that enable rapid optimisation of constructs in BC-producing bacteria. The parts and toolkit described here and in the supplementary materials currently remain untested in other BC-producing strains but we are optimistic that they will work well in all Komagataeibacter and other acetobacter. Indeed modular DNA parts developed previously by us and others for *K. rhaeticus* have been shown to be functional in *Gluconacetobacter xylinus* and *Gluconacetobacter hansenii* (Florea et al., 2016; Gwon et al., 2019; Singh et al., 2020; Teh et al., 2019; Walker et al., 2018). Parts from these past publications, including promoters, RBS sequences and CRISPR-derived gene regulation tools, can presumably be reformatted to work within our KTK system. We look forward to many interesting and innovative genetic parts being added to this toolkit as the growing community in synthetic biology and material sciences begin to produce BC-based ELMs with new functionalities and diverse and exciting properties.

## Materials and Methods

### Strains and culture conditions

The *E. coli* Turbo (NEB) was used throughout this study. Cultures were grown at 37 °C in shaking liquid Lysogeny Broth (LB) (10 g/l Tryptone, 5 g/l Yeast Extract, 5 g/l NaCl) or on LB agar (1% agar), and when appropriate supplemented with ampicillin (100 μg/ml), chloramphenicol (34 μg/ml) or spectinomycin (100 μg/ml). Transformation was done using chemically competent cells.

*K. rhaeticus iGEM* cultures were grown at 30°C in liquid Hestrin–Schramm media (HS) (2% glucose, 10 g/l yeast extract, 10 g/l peptone, 2.7 g/l Na2HPO4 and 1.3 g/l citric acid, pH 5.6–5.8) or on HS agar plates (1.5% agar). When growing shaking cultures the media was supplemented with 2% cellulase (Sigma Alrich, C2730) and, when appropriate, supplemented with chloramphenicol (34 μg/ml) or spectinomycin (100 μg/ml). Electroporation of *K. rhaeticus* strains was performed as described previously (Florea et al., 2016), and transformants screened on 10x Chloramphenicol (340 μg/ml) or 5x spectinomycin HS plates (500 μg/ml). When preparing pellicles for strength tests, *K. rhaeticus* was strains were firstly grown to high density in liquid in the presence of 1x antibiotic. This was then washed twice with HS media, resuspended in fresh cellulose-free HS media to a density of 0.5 OD600. This density was also used as standard for pellicle inoculums. Pellicles were grown stationary at 30°C in fresh cellulase free HS media (75-100 ml), supplemented with antibiotic, and when appropriate 50 mM AHL. Thick pellicles were observed and harvested after 2-4 weeks.

### Molecular techniques, primers and plasmids

Modular DNA parts and plasmids used and constructed in this study are listed in Supplementary Table S1 and S2. Standard PCR or primer joining was used to generate the DNA fragments for the entry-level parts. PCR was performed by standard protocol using a high-fidelity polymerase as per manufacturer’s instructions. Primers are designed to include BsmBI and BsaI required for cloning as detailed in sequences shown in Table G1 and Figure G1. If Entry-level parts were small (<60 bp), instead of a PCR reaction, a primer joining protocol was used. Each primer would cover the target sequence and including the overhang and RE sites detailed in Table G1 and Figure G1. In order to allow primers to anneal together a sequence of overlap of at least 15 bp is required. Oligos (100 μM) are separately phosphorylated with T4 PNK (NEB) before combined and heated to 96°C for 6 min. Samples are annealed by ramping down to 0.1°C per second till 23°C. 10 μl of the mixture was used to clone into the Entry-Level Backbone vectors.

Golden Gate (GG) assembly followed standard GG protocols published (Lee et al., 2015). Briefly T4 DNA Ligase (Promega, C1263), T4 ligase (0.5 μl; NEB, M0202), Type IIs RE (0.5 μl, NEB), vector backbone and inserts were combined to make 10 μl. For optimum results a ratio of insert to backbone of 1:2 was used, *ie*. 50 fmol/μl per insert to 25 fmol/μl of backbone. Thermocyclers were used for assembly with 25 cycles of digestion at 42 °C (2 min) and ligation (16°C, 5 min), before two heat inactivation steps at 60°C (10 min) and 60°C (10 min). After which DNA was be transformed as per standard protocols and plated on prepared selectable plates.

Although GG was used primarily, Gibson cloning was required to adapt more complex entry-level parts, specifically the removal of BsaI restriction enzyme site from the CsgB gene in the synthetic Curli operon, and when building the CsgE-CsgG operon. All Entry-level vectors were sequenced and subsequent plasmids were confirmed with overlapping PCR to ensure all the fragments were present and had combined correctly.

### AHL induction assays

For protein expression assays both *E. coli* and *K. rhaeticus* cultures were grown shaking to mid exponential phase. Cells treated with 50 nM (*E. coli*) or 50 μM (*K. rhaeticus*) of AHL (N-Acyl homoserine lactone, Sigma-Aldridge K3255) and induced for approximately 4 doubling times (i.e. 2 hrs or 16 hrs respectively). Cells were harvested and corrected for by OD_600_ before further analysis performed.

### Chromoprotein expression

*K. rhaeticus* starter cultures were grown in HS-glucose media with chloramphenicol antibiotic at 35 μg/mL for 3 days prior to inoculation of 5 mL of HS-glucose in 24-well deep well plates. The plate was grown static for 5 days at 30°C to grow pellicles. Pellicles were then removed, washed in sterile PBS for 10 minutes and imaged. Cell pellets were obtained by digestion of pellicles with 2% sterile cellulase in PBS (37°C, 250 rpm for 8 hour) before centrifugation at 13,000 rpm and removal of supernatant. Drying of pellicles was done by placing fresh pellicles on open petri dishes and incubating at either 30°C (16 hours) or 60°C (3 hours) or by compressing the pellicle and leaving it a room temperature (24 hours).

### SDS-PAGE and Western Blot

Amyloid associated samples intended for SDS-PAGE analysis were prepared as described previously (Rouse et al., 2017). Briefly, cell pellets were resuspended in 100 μl dH2O and lyophilized. Amyloid containing samples to be monomerised were treated with Formic Acid, frozen and lyophilized a second time. All samples were resuspended in 8M Urea and loading buffer (2% SDS, 0.2 M Tris-HCl pH 6.8, 0.01% bromophenol blue, 10% glycerol, 2 mM DTT) before heated at 95 °C and separated on in 12% SDS-PAGE gels. Proteins were transferred by semi-dry Western Blot onto PVDF membranes visualised with polyclonal antibody and BCIP/NBT (Promega, S3771).

Non-amyloidal samples were treated with protease inhibitor (Roche complete) before fractions separated and harvested. Cell fractions were resuspended in loading buffer and heated at 95°C for 10 min. Supernatant fractions were TCA precipitated, washed with acetone, resuspended in loading buffer and heated at 95°C. Proteins samples were separated on 12-15% SDS-PAGE gels. Gels were stained with SimplyBlue™ SafeStain (ThermoFisher Scientific) and when required, transferred by semi-dry Western Blot onto PVDF membranes visualised with polyclonal antibody and BCIP/NBT (Promega, S3771). Anti-bodies used in this study included a monoclonal α-his-antibody (BioLegend, 652502) and polyclonal α-CsgA-antibody and α-TasA-antibody.

### Transmission electron microscopy (TEM)

*E. coli* and *K. rhaeticus* cultures were prepared with 100 nM AHL but further prepared as described above. Sample Pellets were resuspended in 300 μl HEPES buffer (pH 7.5). For *K. rhaeticus* samples were further treated with 20% cellulase for 4 hrs at 37 °C, before centrifuged again to wash away excess cellulose and resuspended in 300 μl HEPES buffer (pH 7.5). 2 μl samples were spotted onto freshly glow discharged formvar/Carbon on 300 Mesh Nickel grids (Agar Scientific) and blocked for 10 min with BHN1 (1% BSA, 50 mM HEPES, 150 mM NaCl, pH 7.5). Grids were then incubated for 30 min with polyclonal α-CsgA-antibody (1:10 in BHN1, washed twice with BHN2 (0.1% BSA, 50 mM HEPES, 150 mM NaCl, pH 7.5), incubated with secondary anti-Rabbit IgG-Gold coupled antibody (1/10 in BHN1) (Anti-Mouse IgG (whole molecule)–Gold antibody produced in goat (Sigma, G7652), washed twice in BHN2 and HN (50 mM HEPES, 150 mM NaCl, pH 7.5) before stained with uranyl acetate. The FEI Tecnai G2 Spririt TWIN was used to visualise the cells on the grid.

### Material Strength Tests

Pellicles were dried flat using a heated press set to 120°C and 1 ton of pressure. Dog bone test specimens were cut out of the dried cellulose and specimen ends reinforced with card. Dots were marked on the surface of each specimen for the optical measurement of displacement. Tensile tests were conducted with a Deben Microtest Tensile Stage. Seven samples were measured per material, with results normalised for sample thickness.

### Beta-lactamase Nitrocefin Assay

Beta-lactamase activity was measured using Nitrocefin (Stratech, B6052) as per manufacturer’s instructions. Samples were induced as described above, the extracellular supernatant fraction removed post centrifugation. Extracellular fractions were equilibrated and diluted in 10x PBS (pH 7,5) at either 1:100 or 1:10 for *E. coli* or *K. rhaeticus* respectively. Beta-lactamase converts nitrocefin from a yellow to a red substrate and the enzyme activity was calculated as the rate of change of absorbance (490 nm, Synergy HT plate reader) over the linear region of a graph.

## Acknowledgements

We are grateful to A. Kan and N. Joshi for the synthetic Curli template, and to the Chapman Lab and Stanley-Wall Lab for CsgA and TasA-antibodies respectively. The Steve Matthews lab, in particular G. Wu for assisting with Curli protocols and lyophilisation. The Polizzi lab, in particular R. Aw, and both P. Simpson and T. Pape at ICL-EM centre. And finally, to Ellis lab members C. Gilbert and W. Shaw for early discussions on Golden Gate syntax and standards.

