## Supplementary Materials for "Komagataeibacter tool kit (KTK): a modular cloning system for multigene constructs and programmed protein secretion from cellulose producing bacteria"

### Supplementary Figures

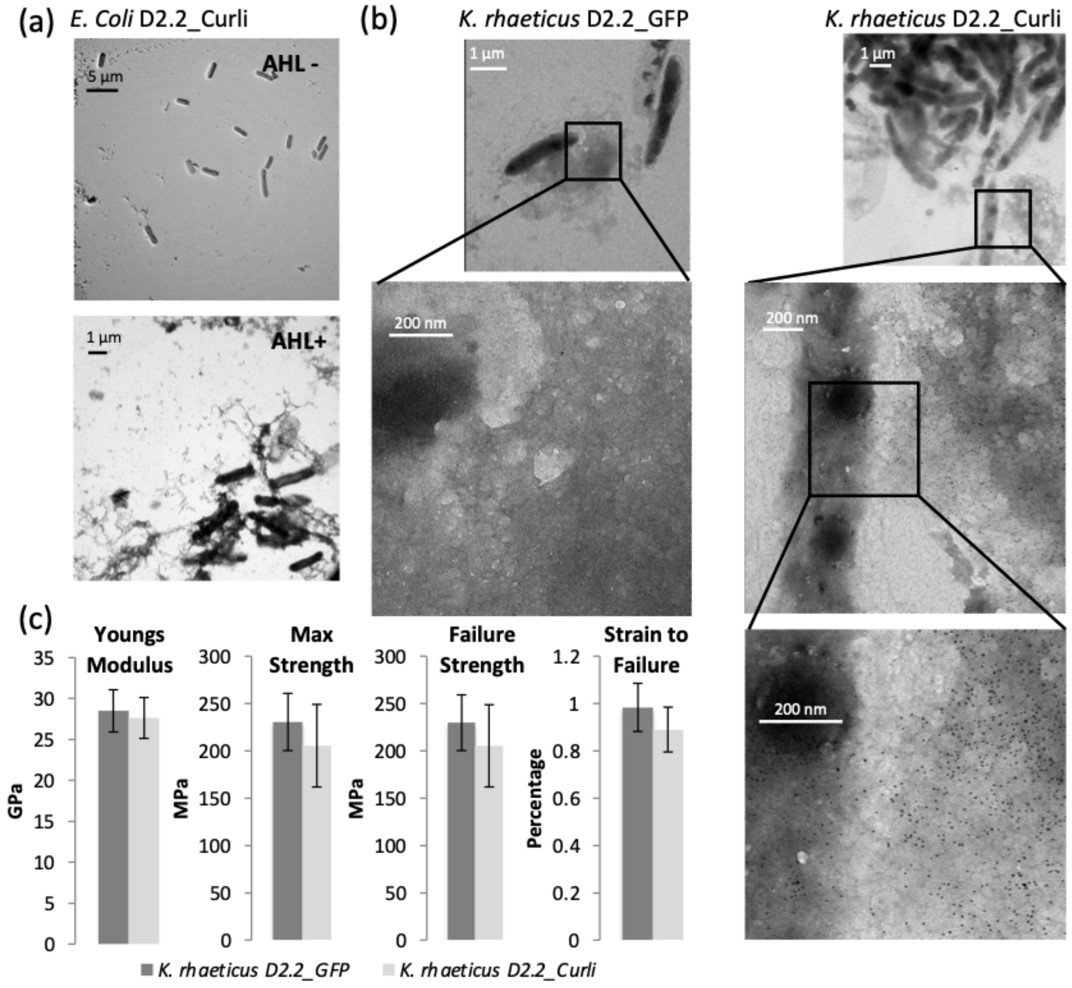

**Supplementary Figure 1. Characterisation of *K. rhaeticus* and *E. coli* heterologouly expressing the whole *E. coli* Curli system (D2.2_Curli).** (a) TEM images of *E. coli* strains with (AHL +) and without (AHL -) AHL induction. (b) Immunogold TEM (α-CsgA and 10 nm anti-Rabbit IgG-Gold coupled antibody) samples are of AHL treated *K. rhaeticus* strains D2.2_Curli or D2.2_GFP (c) Structural pellicle test performed on AHL treated *K. rhaeticus* pellicles. Samples size for each strain n =7

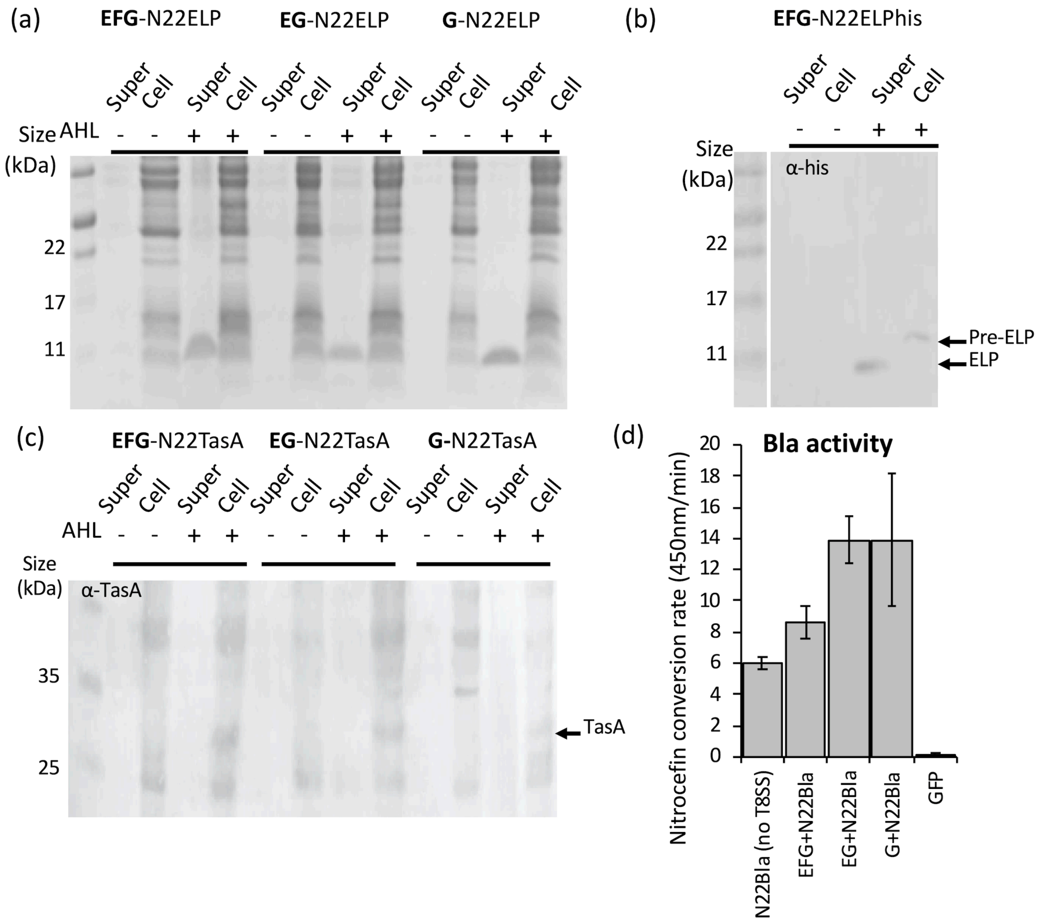

**Supplementary Figure 2. Exploiting the T8SS using KTK to secrete heterologous cargo proteins in *E. coli*.** (a) 15% SDS-PAGE gels of *E. coli* strains expressing various T8SS and N22ELPhis as the cargo protein, both cell fraction and supernatant fraction are shown, samples were induced with AHL (+) or uninduced (-) (b) Western Blot of samples extracted from *E. coli* expressing T8SS including CsgE-CsgF-CsgG and the N22ELPhis protein. ELP-his is detected using α-his antibody. Full length pre-ELP (expected size 12,6 kDa) and secreted and processed ELP (expected size 10.6 kDa) are indicated by arrows. (c) Western Blot of Formic acid treated *E. coli* strains expressing various T8SS and N22TasA, TasA protein (30 kDa) is only observed in cell fractions. (d) Nitrocefin assay with supernatant extracted from induced *E. coli* strains expressing N22Bla with and without T8SS components. D2.2_GFP was used as a negative control (GFP in graph). Error bars represent SD of three replicates. The assay itself was repreated six times in all cases the presence of a T8SS module resulted in significant increased enzyme activity in the supernatant.

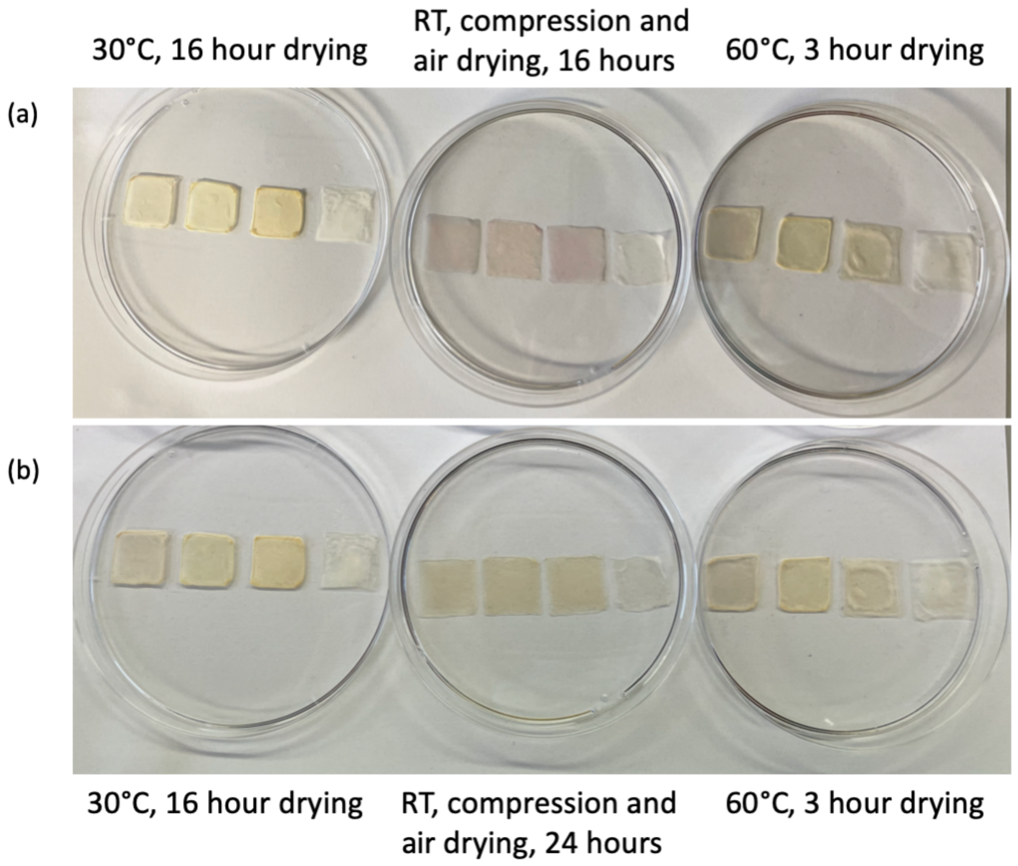

**Supplementary Figure 3. Dehydration of chromoprotein pellicle samples** (a) Pellicle squares (left to right) grown from bacteria with blue, red and purple chromoprotein expression and wildtype cells, were imaged 16 hours after being subjected to one of 3 dehydration methods: open dish incubated at 30°C in oven with fan, compression followed by open dish at room temperature, open dish incubated at 60°C in oven with fan for 3 hours, then stored for the remaining 13 hours. (b) The same pellicles as in (a) after a further 8 hours incubation at room temperature, showing that the completed dehydration of all samples led to loss of colour.

### Supplementary Tables.

**Table S1. KTK library. (excel sheet)**

**Table S2. Plasmids used in this study (excel sheet)**

**Table S3. Detailed KTK overhang sequences (excel sheet)**

### KTK: a detailed introduction and guide

The Golden gate (GG)-based KTK system is best described in cloning levels. It starts with a number of individual Entry-level plasmids, each contain a single promoter, RBS, CDS regions or terminator. In Level 1 cloning entails combining Entry-level Parts with a Destination vector backbone to form a single full genetic Cassette or Transcription unit (TU). Level 2 cloning allows for the combination of multiple TU using Destination 2 vector backbones. The resulting plasmids can be once again combined in a 3rd cloning level to form plasmids with cassettes in multiples of 2, Fig G1 and Table G2.

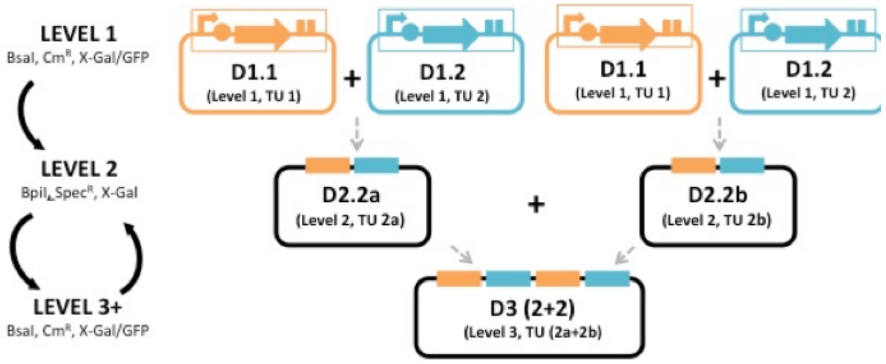

**Supplementary Fig G1**. Golden Gate cloning schematic of KTK. The KTK kit a GG-based cloning system best described in levels. Level 1 and 3 make use of BsaI, selection on Chloramphenicol and counter-selection using X-gal and GFP. Level 2 makes use of BpiI, selection on Spectinomycin and counter-selection using X-gal screening. Level 1 involves combining the Entry-level Parts to form single TU. There are two types of Destination Level 1 Cloning vector backbones resulting in D1.1 and D1.2 plasmids. In a similar fashion the single transcription units can then be combined in pairs in a second cloning level, and in third cloning level allows for the combination of four constructs. The Level 3 plasmid behaves similarly to the D1.1 plasmids and has overhangs that allow for further iterations of cloning if required. Hence 3+ cloning rounds are possible.

#### Entry-level Parts

In order to translate a sequence a complete genetic cassette or transcription unit (TU) is required. In Golden Gate (GG) cloning these TU are divided into Parts: a Promoter (E1.1), RBS (E1.2), Coding sequence (CDS) (E1.3) and a Terminator (E1.4) Part. (Fig G2a). A collection of Entry-level parts, the KTK library, is currently being expanded on and will be made available via Addgene.org in the near future. This modularity can enable you to quickly generate and test a library of TU that containing variable Parts. The kit also allows for the introduction two-Part fusion proteins (separate C- or N-terminal domain to the CDS) enabling one to generate signal peptides, tags (e.g. His-tag) or more complex protein fusion (e.g. GFP) (Fig G2a). The sequence of each Part is cloned separately into an Entry-level vector backbone, KTK_001, replacing the GFP cassette (Fig G2b). KTK_001 is an ampicillin-resistant *E. coli*-based high-copy number plasmid that does not have a compatible origin of replication for Komagataeibacter species and is unable to replicate in the host strain.

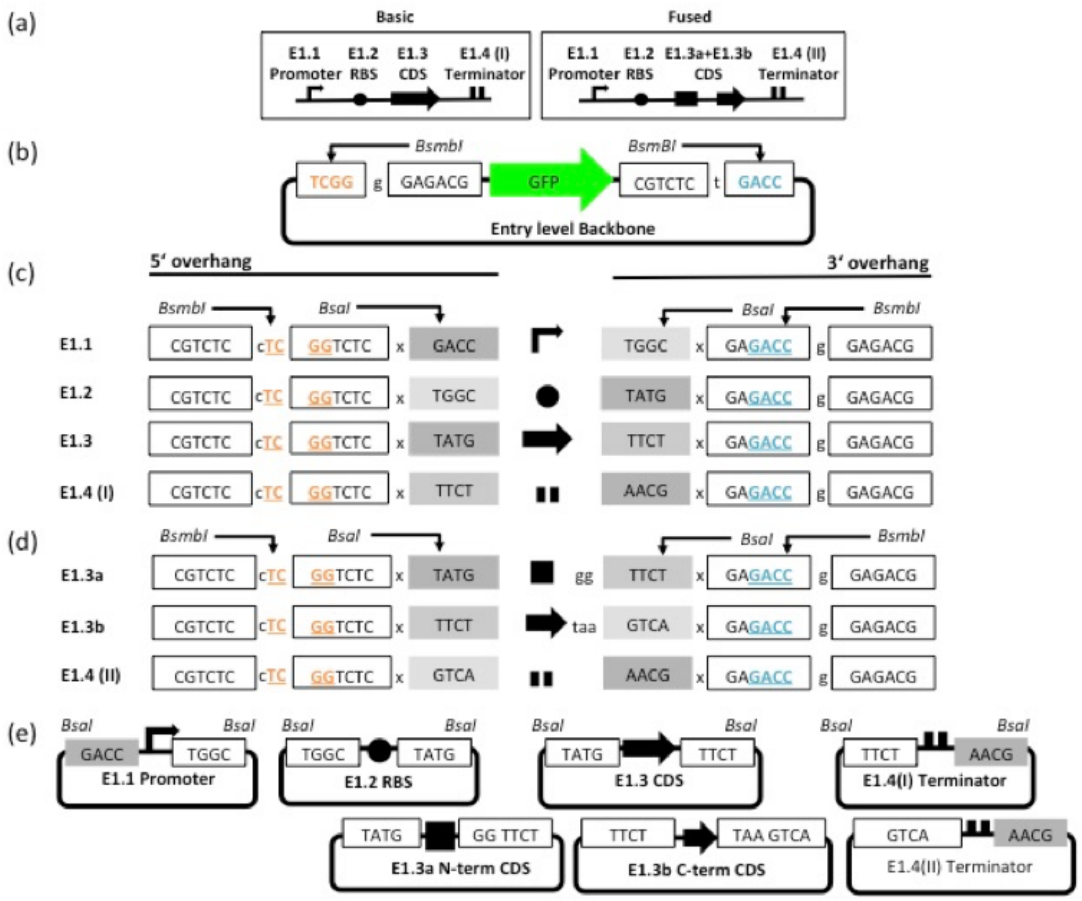

**Supplementary Figure G2** Detailed Entry-Level cloning. (a) Illustration of Basic and fused Transcription units (TU). TU are made up of Promoters (angled arrows), ribosome bindings sites (circles), CDS regions (boxed arrows), and a terminator (paralleled lines). (b) The Backbone Entry-level vector, KTK_001, has a GFP expressing cassette flanked by BsmBI RE sites that generate complementary overhangs to the Entry-Level DNA sequences (shown in orange and teal). These facilitate insertion of the individual part. (c) 5' and 3' sequences for standard Entry-level parts are generated by primer overlap assembly or PCR, and are made with essential 5' and 3' flanking sequences. These flanking sequences include type IIS RE sites that will result in complementary GG overhangs. The sequences for BsmBI and BsaI type IIS RE sites are given in boxes and black arrows extending from these indicate position where the enzymes cut and the overhangs that are created. BsmBI overhangs are shown in orange and teal. The overhangs generated by BsaI are shown in grey boxes. (d) 5' and 3' sequences for fused-CDS domains. (e) The seven different kinds of Entry-level Plasmid Parts.

#### Basic Part Vectors

Each of the four basic Parts to a TU has a unique overhang sequences (Fig G2c). The individual parts are created by PCR or Primer joining and are designed to have specific GG-enabling overhangs, BsmBI and BsaI restriction enzyme (RE) sites (Fig G2c, Table G1). The BsmBI site facilitates insertion of individual Parts into the Entry-level Vector backbones and the resulting vectors are then screened on ampicillin ($Amp^R$), where the loss of GFP implies the correct insertion of the Part (Fig G2e). Entry-level Part vectors need to be sequenced verified before continuing with the next step (Table G3). Consequent steps do not require re-sequencing and vectors can be checked by RE digest/overlapping PCR alone.

However, each individual project will require a unique Entry-Level Part E1.3, corresponding to the specific CDS required. When designing the PCR for the E1.3 Part it is important to note that 5' overhang generated by the Primer sequence (Fig 2c) already includes a start sequence, however the 3' region does not include a stop codon. Therefore, depending on your downstream application you may need to include a stop codon, but not a start codon in your primer design (this is not the case for fusion-proteins). Importantly, none of the Parts used in any KTK studies should include sequences that correspond the BsmBI, BsaI or BpiI/BbsI RE sites, as this will prevent downstream GG cloning. If these sequences do exist we recommend removing them, e.g. with overlap extension PCR, with internal primers designed to safely mutate the site.

#### C- or N-terminal CDS-fusions

When making a TU with a fusion protein the general protocol is similar to what was described for the Basic Entry-level Part. However, there is an important difference with regard to the overhangs used to make the CDS Entry-level Parts (E1.3a and E1.3b) and the Terminator (1.4(II)) (Fig G2d) If you intend to make a C-terminal tag, e.g. a Signal Peptide (SP), this would mean that the SP sequences is cloned as a E1.3a Part the CDS into E1.3b Part. In the case of a N-terminal Tag, e.g. a His-tag, this would mean the CDS is cloned into an E1.3a Part and the His-tag into E1.3b Part. Importantly the

CDS E1.3a sequence should not include a stop codon to ensure a continuous ORF with the E1.3b sequence.

In order to ensure the overhangs created by the RE do not introduce a frame shift mutation, a few linker base pairs between Parts E1.3a and E1.3b have been introduced into the designed overhang sequence (Fig G2d). These extra base pairs result in a linker region that when translated encodes glycine and serine amino acids.

**Level One Cloning: Creating Transcriptional Units.**

There are a four Level 1 backbone Destination vectors (Fig G3). The differences are associated with counter selectable marker (X-gal or GFP) and overhangs available for Level 2 cloning. Each backbone allows for combination of the Entry-level Parts to form single TU. Therefore, if you are making a simple one level genetic cassette, it does not make a difference which of the three you use. However, if intend to combine cassettes in Level 2 and onwards, the specific destination vector chosen does make a difference with regards to the order in which cassettes can be combined (Fig G1). Further, if you intend to express a GFP-based protein in the desired construct, then the X-gal counter-selectable marker (LacZ) needs to be used.

Constructs are built by combining a Destination Vectors backbone with the Entry level Parts in a GG reacting based on BsaI RE activity. Clones are selected for on chloramphenicol ($Cm^R$) and for the absence of GFP or LacZ activity (blue/white selection on X-gal media).

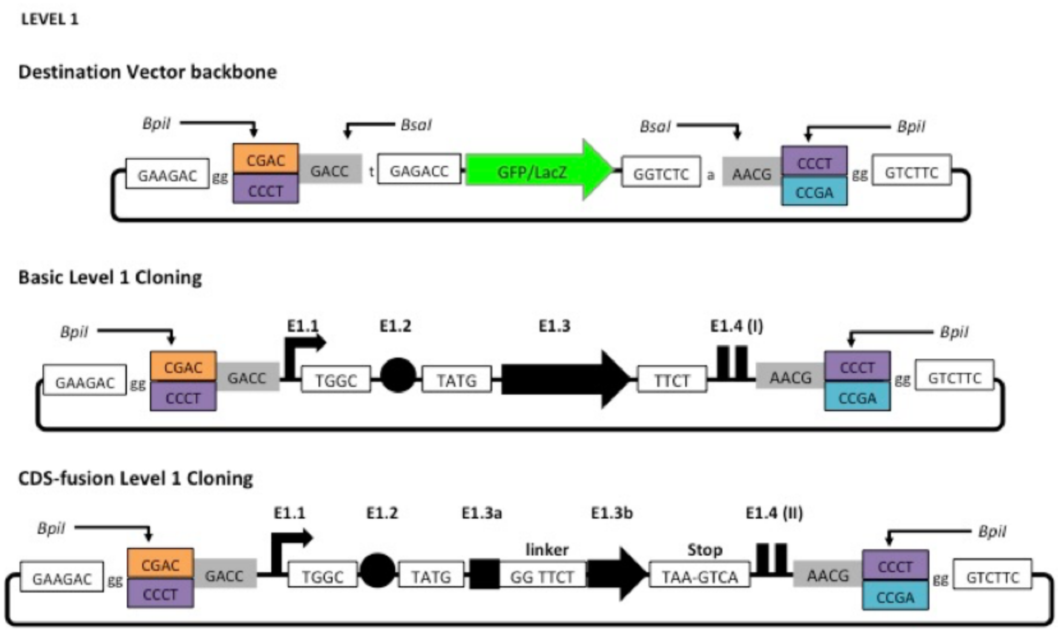

**Supplementary Figure G3.** Detailed visualisation of Level 1 KTK Cloning. The Destination Vector backbones with variation in counter-selectable marker (GFP/X-gal) and the overhangs used to facilitate GG-cloning. Type IIS RE sites are given in 6-lettered boxes and black arrows extending from these indicate position where the enzymes cut and the overhangs that are created. The overhangs used in Level 1 cloning are shown in grey/white boxes. And the overhangs required for Level 2 cloning are shown in blue/purple/orange. TU are made up of Promoters (angled arrows), ribosome bindings sites (circles), CDS regions (boxed arrows), and a terminator (paralleled lines).

#### Level two and three cloning: making multi-cassette-containing plasmids

The DNA sequences flanking the Type IIS RE-sites created in Level 1 allow for the GG-based cloning of two TUs into one of the two Destination 2 Backbone vectors (Fig G1 and Fig G4) using BpiI digestion and ligation. These vectors have Spectinomycin resistance and can be screened for the absence of LacZ (i.e. white on X-gal) to determine successfully cloning. In Level 3 cloning the plasmids generated in Level 2 can be combined again into specially designed vector backbones. Here BsaI is used to generate the overhangs and once again correct white clones are selected for on spectinomycin and X-gal media to show loss of lacZ from the vector. The vector backbones used to make the Level 3 (2+2) plasmids are technically similar to those used to make D1.1 plasmids at the 1$^{st}$ cloning level. Therefore, theoretically users can iterate this schematic (Fig G1) and continue to build more complex cassettes and plasmids if required, alternating between BpiI- and BsaI-based Golden Gate assembly reactions. If the construct you are building is made up of an odd number of modules, spacer cassettes available in the kit that can act as intergenic regions.

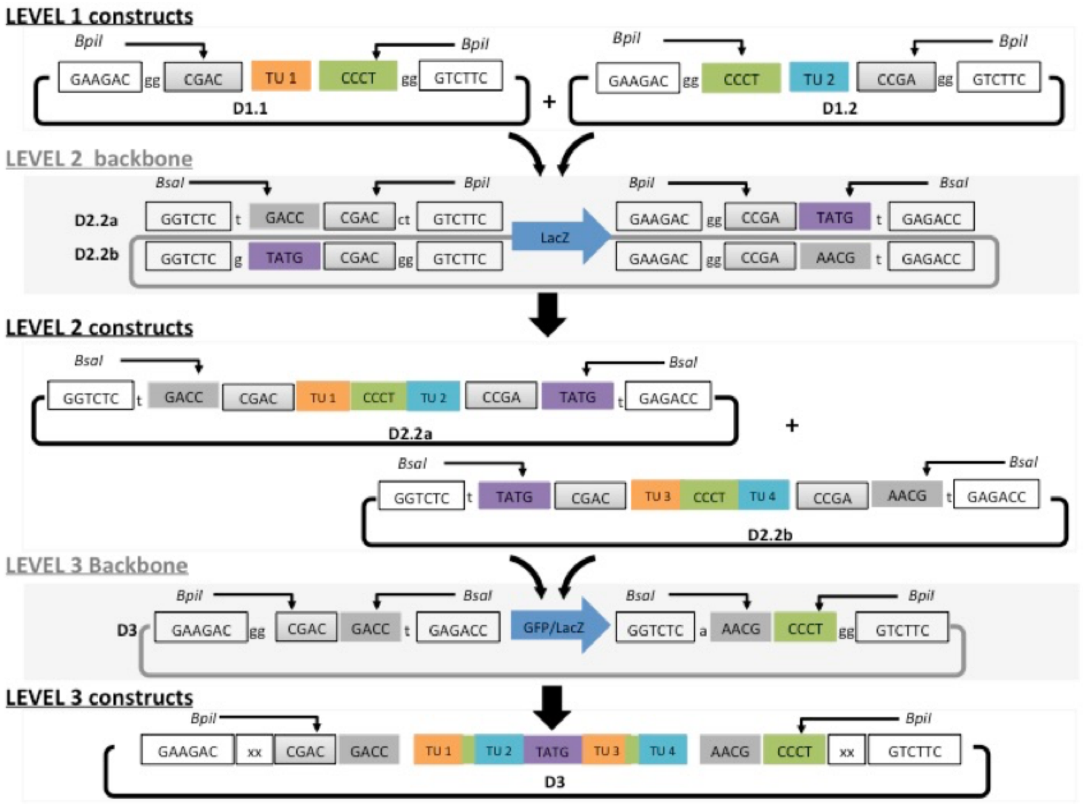

**Supplementary Figure G4.** Detailed schematic of Level 2 and level 3 cloning. Type IIS RE sites are given in 6-lettered boxes and black arrows extending from these indicate position where the enzymes cut and the overhangs that are created. The overhangs in GG cloning reactions are shown in grey/green/purple. The Backbone vectors are indicated in grey with there are two possible Level 2 destination vector backbones (DD2.2a and D2.2b) and one Level 3 (D3). The resultant constructs are shown in black. TU have been numbered and colour coded in orange and blue.

#### Table G1 Entry Level Overhang sequences

| Backbone = KTK_001 | Type of entry | 5'OH sequence | 3'OH sequence (not primer i.e. should not be reverse complemented) | Selection of clone | RE |
|---|---|---|---|---|---|
| | | **Basic Part Vector** | | | |
| Entr 1.1 | promoter | CGTCTCCTCGGTCTC T GACC | TGGC TGAGACCGGAGACG | Amp$^R$ & loss of GFP | BsmBI |
| Entr 1.2 | RBS | CGTCTCCTCGGTCTC A TGGC | TATG TGAGACCGGAGACG | Amp$^R$ & loss of GFP | BsmBI |
| Entr 1.3* | CDS | CGTCTCCTCGGTCTC C TATG | TTCT TGAGACCGGAGACG | Amp$^R$ & loss of GFP | BsmBI |
| Entr 1.4(I) | Terminator | CGTCTCCTCGGTCTC T TTCT | AACG TGAGACCGGAGACG | Amp$^R$ & loss of GFP | BsmBI |
| | | **Fusion-Protein Part Vector** | | | |
| Entr 1.3a** | N-term. fusion | CGTCTCCTCGGTCTC C TATG | GG TTCT T GAGACCGGAGACG | Amp$^R$ & loss of GFP | BsmBI |
| Entr 1.3b* | C-term. fusion | CGTCTCCTCGGTCTC A TTCT | TAAGTCA A GAGACCCGAGACG | Amp$^R$ & loss of GFP | BsmBI |
| Entr 1.4(II) | Terminator- | CGTCTCCTCGGTCTC A GTCA | AACG T GAGACCCGAGACG | Amp$^R$ & loss of GFP | BsmBI |

5' and 3' overhang sequence detailed. BsmB1 underlined in blue, BsaI site underlined in orange. Complementary Golden Gate overhangs are coloured in complementary colours. *Inserts into this entry vector require a Stop codon ** Insert into this entry vector cannot have a Stop codon.

**Table G2 Level 1-3: Cloning schematic**

| Backbone vector | Part used | Result | Selection | Type IIs RE |
|---|---|---|---|---|
| **Level 1** | | | | |
| KTK_009 /KTK_136 | E1.1 + E1.2 + E1.3 + E1.4 (I) | KTK_D1.1 | $Cm^R$ & X-gal/GFP | BsaI |
| KTK_010/KTK_137 | OR | KTK_D1.2 | $Cm^R$ & X-gal/GFP | BsaI |
| KTK_011/KTK_138 | E1.1 + E1.2 + E1.3a + E1.3b + E1.4(II) | KTK_D1.3 | $Cm^R$ & X-gal/GFP | BsaI |
| **Level 2** | | | | |
| KTK_015 | D1.1 + D1.2 | KTK_D2.2a | $Spec^R$ & X-gal | BpiI/(BbsI) |
| KTK_016 | D1.1 + D1.2 | KTK_D2.2b | $Spec^R$ & X-gal | BpiI/(BbsI) |
| **Level 3** | | | | |
| KTK_009 /KTK_136 | | KTK_D3.1 | $Cm^R$ & X-gal/GFP | BsaI |
| KTK_010/KTK_137 | D2.2a + D2.2b | KTK_D3.2 | $Cm^R$ & X-gal/GFP | BsaI |
| KTK_011/KTK_138 | | KTK_D3.3 | $Cm^R$ & X-gal/GFP | BsaI |

**Table G3 Useful Primers**

| Primer Name | Primer use | |
|---|---|---|
| KTK_seq_FP | Sequencing primer for Entry level cloning flanking left of the insertion | TATAGTCCTGTCGGGTTTCGCC |
| KTK_Seq_RP | Sequencing primer for Entry level cloning flanking right of the insertion | CCGGTGAGCGTGGGTCCCGCGGTATC |
| KTKStdFrdD1_D2 | Primer for region left of insertion in KTK_D1 and KTK_D2 constructs | AGGGCGGCGGATTTGTCC |
| PQE30Rev | Primer for region right of insertion in KTK_D1 and KTK_D2 constructs | GTTCTGAGGTCATTACTGG |
